# Identification and evolutionary analysis of a Triticeae tribe specific novel non-autonomous DNA transposon in *DREB* related *Dehydration Responsive Factor1* gene

**DOI:** 10.1101/2022.03.05.482545

**Authors:** K Thiyagarajan, A Latini, C Cantale, E Porceddu, P Galeffi

**Affiliations:** Sant’ Anna School of Advanced Studies, Piazza Martiri della Libertà, 33, 56127 Pisa, Italy; Casaccia Research Centre, ENEA (Italian National Agency for New Technologies, Energy and Sustainable Economic Development), Via Anguillarese 301, 00123 Rome, Italy

## Abstract

A non-autonomous DNA transposon was identified in the *DRF1* gene, belonging to the *DREB* gene family, the presence of this element was initially assessed in the *Triticum durum DRF1* gene and subsequently it was also identified in *Aegilops speltoides* and *Triticum urartu DRF1* genes. The DRF1 gene consists of four exons and three introns, the transposon carrying core element is inserted between the first and the third introns. Our studies identified inverted repeats, target site duplications and the presence of many internal reverse and direct short tandem and long tandem repeats, that all represent signals of a transposable element. Based on transposon specific sequence and position of the terminal inverted repeats, a possible transposition mechanism was inferred. As the identified transposable element does not possess a sequence coding for a transposase enzyme, it represents a non-autonomous element. The transposon encompasses a core element with two small, transcribed regions (Exon 2 and Exon 3) that are combined by alternative splicing during gene expression and an intron (intron2). A possible role of this non-autonomous DNA transposon in the alternative splicing regulation was investigated by a genomics approach. Divergence time analysis supported the relatively recent evolution of this transposon in Triticeae comparing to other tribes and further there is no footprints or highly disrupted footprints sequence such as TIR, TSD in other earlier evolved Poaceae member species were observed, which revealed the novelty and well-preserved nature of these signals in Triticeae. While other monocots (apart from Poaceae) and dicots, including *Arabidopsis thaliana*, neither showed this transposon insertion and nor revealed the existence of alternative spliced gene transcripts. In Poaceae members the core element is well preserved with disturbed transposon and transposon signals, while the tribe Triticeae especially wheat, its progenitors have intact DRF1 transposon and its signals.

## Introduction

### Transposable Elements

Transposable Elements (TE) consist of repeated sequences of DNA with defined structures that can move from one region to another directly (through DNA) or indirectly (through RNA intermediate, so-called retro elements). Transposase makes a staggered cut at the target site producing sticky ends, cuts out the transposon and ligates it into the target site. A DNA polymerase fills in the resulting gaps from the sticky ends and DNA ligase closes the sugar-phosphate backbone. This mechanism of insertion produces site duplication in the target, or the insertion sites of DNA called Target Site Duplication (TSD), a key signature of DNA transposon with Terminal Inverted Repeat (TIR). The transposons may be identified by short direct repeats (a staggered cut in the target DNA filled by DNA polymerase) followed by inverted repeats (which are important for the transposon excision by transposase) (Craig, 1995). Depending on their transposition mobility transposons are typed into autonomous, which are transposed by themselves through *cis* mediated process, since they encode an active transposase, and non-autonomous elements, which depends on cognate autonomous elements for movement, due to their inability to code transposase. The full-length DNA transposons are hypothesized to be the evolutionary progenitors of MITEs, based on sequence conservation of miniature inverted-repeat transposable elements (MITEs) with autonomous transposon terminal inverted repeats (TIRs) and target site duplications (TSDs) (Bureau, 1992). MITES do not have internal homology to their parental autonomous transposons and their internal regions are often non homologous. MITEs have been amplified to a high copy number in several plant genomes, as in rice, which contains more than 90,000 MITEs grouped into approximately 100 different families (Jiang *et al*., 2004). Repeats have precise relevancy to transposons; repeated sequences have also been also isolated from Triticeae group, such as *Aegilops* (Rayburn and Gill 1986; Salina *et al*., 2006). Transposable elements were previously considered as selfish parasites (Orgel and Crick 1980), though these elements causing mutations by chance, but vastly influencing genome structural and biological organization (Kidwell and Lisch 2001). Besides, the transposable elements would be activated more during the stress conditions of the host (Capy et al. 2000), which allows an organism susceptible to have the changes on its hereditary material through the mutual relationship between the host genome and the transposable element rather than the parasitic and tends to have an evolutionary role in stress conditions by acting as “natural genetic engineers” (Shapiro 1999). *DRF1* (*Dehydration Responsive Factor 1*) gene belongs to DREB gene family, where the non-autonomous DNA transposon identified in this study is inserted. Genes in this family are generally present in several copies (Liu et al 2018) either as a whole sequence or as a partial one (the core portion), possibly due to a transposon mediated mechanism. In this article we have described clearly about the identification, organization and evolutionary importance of this transposon in DRF1 gene in monocots particularly in poaceae and postulated a hypothesis about the absence of this transposon in DREB genes of dicots.

## Material and Methods

### Sequences

The sequences of *Aegilops speltoides DRF1* gene, namely FJ858188, FJ858187 and FJ843102, were used in the analysis. The sequence of *Triticum durum DRF1* gene (EU089819) was also analysed and compared with the above ones. Additional gene sequences were retrieved from Genbank: *T. durum* (JN571425), *A. tauschii* (EU197052), *A. columnaris* (GU017675), *P. juncea* (JF766085), *L. chinensis* (JF754585), *H. vulgare* (AY223807). Furthermore, sequences (*T. aestivum* Chr1A, positions-<u>391258134-391259647</u>, *P. halli* V3.2 (positions: Chr:03, 25976840-25978153, *P. virgatum* v5.1 (Chr03N, positions: 34762702-34764024 *O. brachyantha* Chr5 V1.4b, positions: 9533878 -9535170, *O. glabberima* V1 Chr5, positions: 12071071 to 12072366, *O. barthi* V1 Chr:5, positions: 12325233 to 12326528, *S. italica* V2, Chr:25994256 to 25995588, *B. distachyon* Chr2, V3, positions: 29255734 to 29256748, *P. virgatum* V5.1 Chr03N, positions: 34762702, *Z. mays* Zm-B73-REFERENCE-NAM-5.0, Chr8, positions: 96775872 to 96777269) were retrieved from public genome databases such as https://phytozome-next.jgi.doe.gov/blast-searchhttps://plants.ensembl.org/Multi/Tools/Blast

### Computational tools

Repeats, such as terminal inverted repeats, palindromes and other repeats, were identified by computational tools. EINVERTED (EMBOSS) (http://emboss.sourceforge.net/apps/cvs/emboss/apps/einverted.html), Palindrome (EMBOSS) (http://emboss.sourceforge.net/apps/cvs/emboss/apps/palindrome.html), REPuter (Kurtz *et al*., 2001) (http://bibiserv.techfak.uni-bielefeld.de/reputer/submission.html) and Tandem Repeat Finder (Benson, 1999) (http://tandem.bu.edu/trf/trf.html) servers were used. Different lengths of repeats were identified adjusting the visualization window at REPuter server. A secondary DNA structure was predicted using MFOLD (http://www.unafold.org/mfold/applications/dna-folding-form.php). CpGplot (EMBOSS) was used to identify and plot CpG islands in *DRF1* gene sequence (https://www.bioinformatics.nl/cgi-bin/emboss/cpgplot). CENSOR server (Kohany *et al*., 2006) (http://www.girinst.org/censor/index.php) was used for comparison and annotation of the identified repeats at GIRI database. Blast search against TREP (Triticeae Repeat Sequence Database) (http://wheat.pw.usda.gov/ITMI/Repeats/blastrepeats3.html) was also carried out.

The analysis of the exon intron splice sites and branch points was carried out using NetGene2 (http://www.cbs.dtu.dk/services/NetGene2/) using *Arabidopsis thaliana* as model system. The microsatellites were computationally analyzed by IMEx-web server at http://www.mcr.org.in/imex/, Webserver for extracting microsatellites from genome sequences (Mudunuri and Nagarajaram, 2007). Divergence time tree was calculated using the RelTime method (Tamura et al 2012), relative divergence time was generated using the Maximum Likelihood method and Tamura-Nei model (Tamura and Nei 1993). Totally 21 nucleotide sequences and 1790 positions inside these sequences were analysed using MEGA 11 (Tamura 2021). Initially the transposon was inferred and annotated from *DRF1* gene of *T. durum* and *A. speltoides* and subsequently with other species.

## RESULTS

### A novel non-autonomous DNA Transposon in the *DRF1* gene

A novel DNA transposable element, approximately 1.5 Kb long, was identified by bioinformatics tools between Intron 1 and Intron 3 of the *DRF1* gene (See Fig. 1). Initially, it was identified in *T. durum* and subsequently in *A. speltoides, T. urartu DRF1* gene and was also identified in other Triticeae species. As no sequence for transposase was found in the identified transposon, it resulted to be a non-autonomous transposon. The TIR region covering the transposon part was also in various species/accessions including *T. durum, T. urartu, A. speltoides A. tauschii*, as shown in Fig. 1. The sequence of this new transposon was deposited in Repbase (https://www.girinst.org/server/publ/AsDRF1/ and https://www.girinst.org/2009/vol9/issue2/TdDRF1.html) (Karthikeyan et al., 2009, Thiyagarajan et al., 2009).

**Figure 1.**
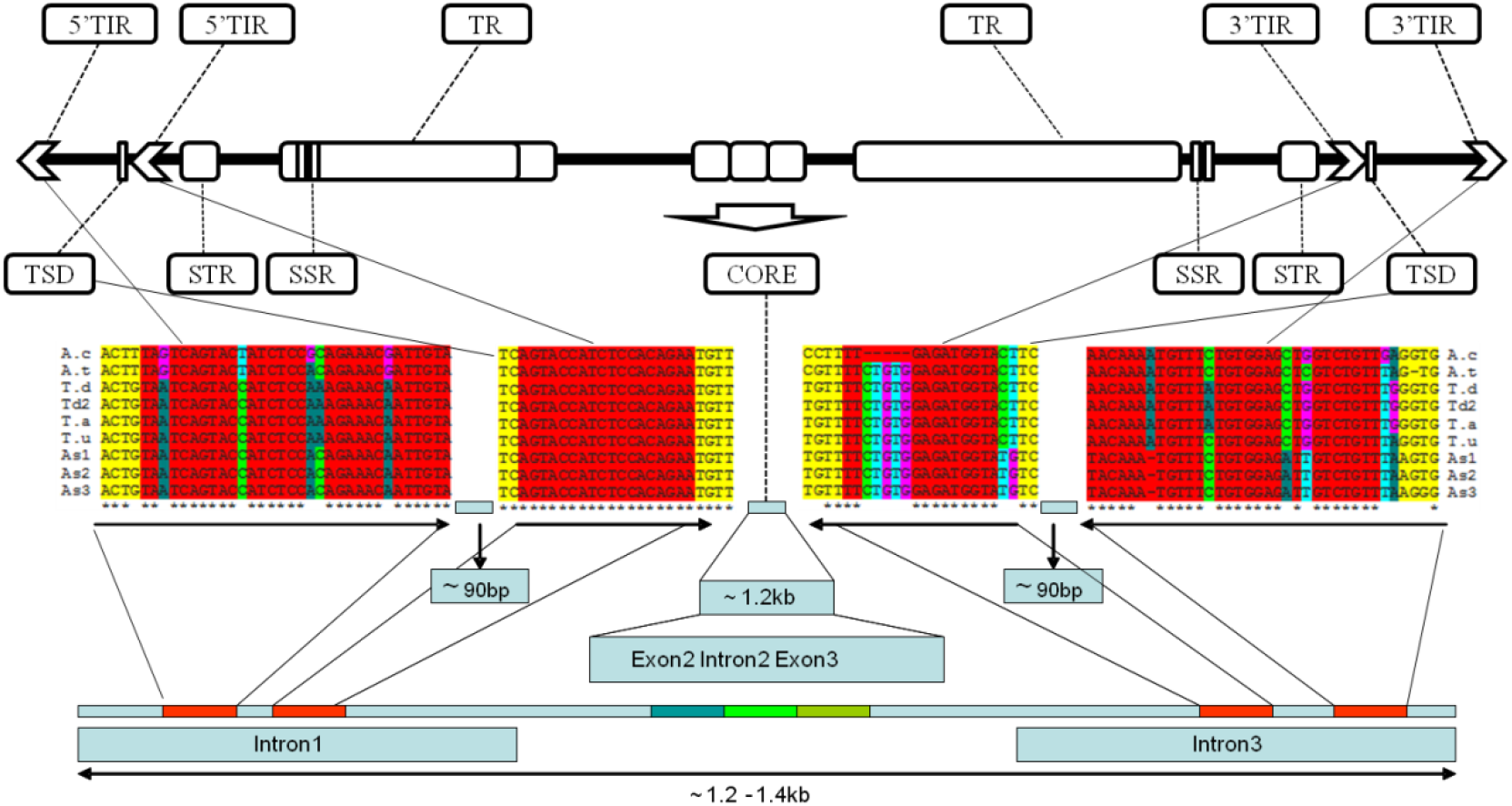
Structure and alignment of 18bp TIRs and 32bp TIRs in *DRF1* genes from different *Triticeae* members. GU017675; *A*.*tauschii*, EU197052; *T*.*durum*; JN571425; *T*.*durum* accession 2; *T*.*aestivum*: sequence from Chr. 1AL at URGI (http://urgi.versailles.inra.fr); *T*.*uratu*: lab sequence acc. 57_7; *A. speltoides* accession 1: FJ843102; *A. speltoides* accession 2: FJ858188; *A. speltoides* FJ858187. TIR-Terminal Inverted Repeat, TR-Tandem Repeats, (STR-Subterminal Repeats, TSD-Target Site Duplication, Core-Exon2, Intron2, Exon3, SSR-Simple Sequence Repeat.

### Computational analyses of Repeats, Palindromes, Tandem, Inverted Repeats inside transposon

Reported results refer to *A. speltoides ssp speltoides* (FJ858188). The direct and reverse repeats were computationally identified by the REPuter server using both default and user-defined values for parameters. Several repeats were observed, ranging from 12 bp to 88 bp. Two 88bp repetitions were observed with high confidence (E-value 1.04e^− 42^, distance -2) starting at 1479 and 1643, respectively. Besides direct repeats, also several reverse repeats were identified, like the 19 bp repeat at 1023 (E-value 1.05e^−05^ and distance 0). In total about 51 repeats were observed in the whole gene of *A*.*speltoides DRF1*, whose 12 were reverse repeats (Fig. 2).

**Figure 2.**
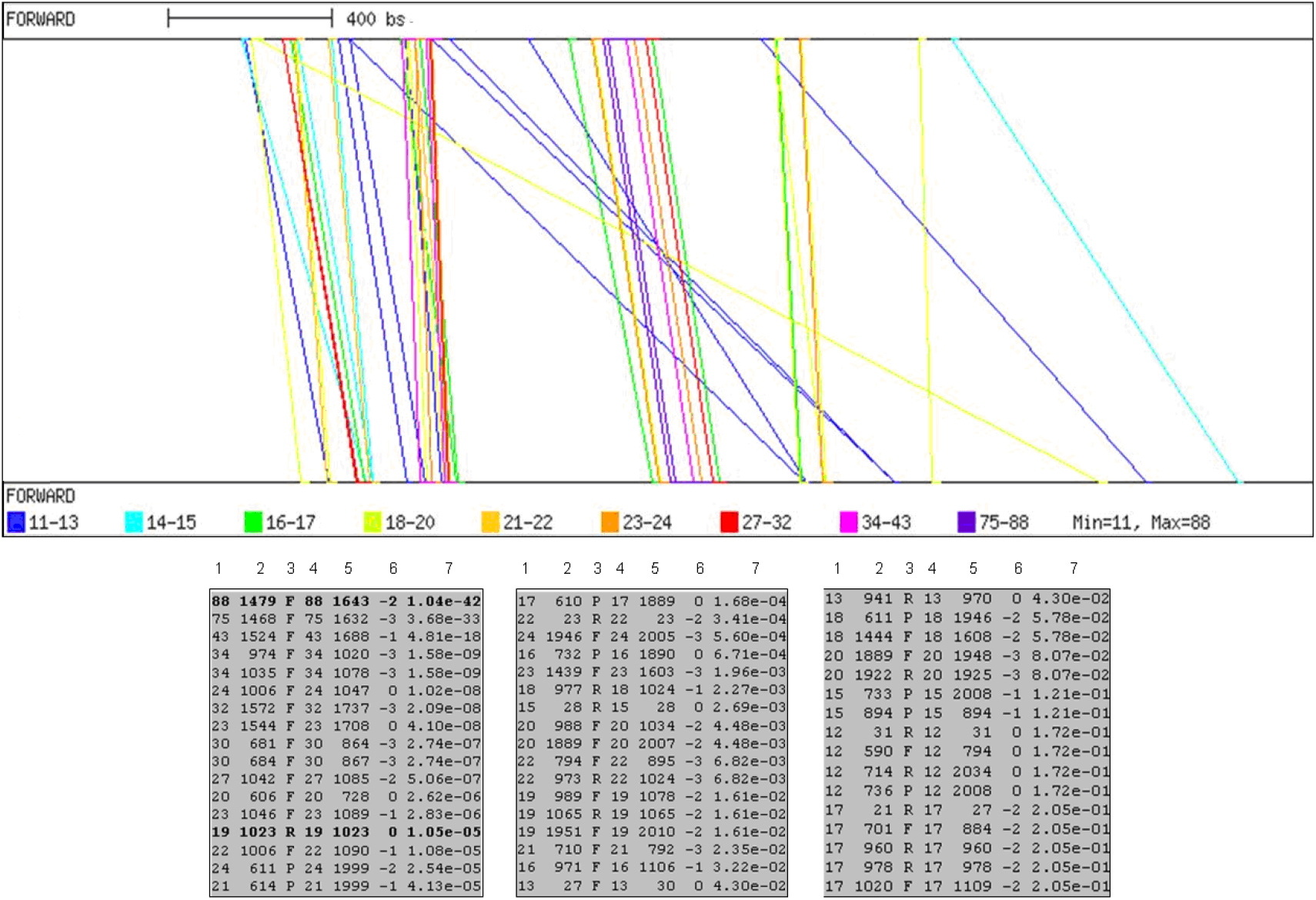
Panel A: REPuter graphical output. All repeats are shown, coloured according to the length. To keep the starting position information visible, each part of a repeat is displayed on a separate strand. Panel B: The tables contain the identified repeats, described by 7 values: columns [1] - length of the first part of the repeat, [2] - starting position of the first part, [3] - match direction, [4] - length of the second part of the repeat, [5] - starting position of the second part, [6] - distance of this repeat, [7] - calculated E-value of the repeat.

Using a window of 8bp for analysing the gene sequence, 8 palindromes, 8bp length, were identified. The same analysis was carried out in *Triticum durum DRF1* gene sequence and 7 palindromes were observed. Three palindromes were shared between the two species, revealing the evolutionary significance of conserved palindromes. The first two palindromes (at 1408 and 2074 in *AsDRF1*) are located within Intron 3, respectively 26 bp after Intron 3 beginning and at the end of Intron 3. The third one is located within Exon 4, 165bp downstream to AP2 domain, in both species. Using Tandem Repeat Finder server, some tandem repeats were found in both species. A long, twice repeated, tandem repeat about 164bp length was identified in *A. speltoides DRF1* gene. It was not present in *T. durum DRF1* gene. Furthermore, a 42bp short tandem repeat was identified in both species. The shared TR was repeated 4.5 times in *Triticum durum DRF1* gene and 3.2 times in *A. speltoides DRF1* gene. Analyses of Inverted repeats carried out using *TdDRF1* gene sequence identified 18bp TIR in *Triticum durum*, located in Intron 1 and Intron 3, characterized by no mismatch. The presence of 2bp TSD (TC), flanking the TIR, and 4bp STR (Sub Terminal Repeat) / ITSD (Internal Target Site Duplication) sequences (TGTT) were manually assessed. An identical 18bp TIR sequence was manually identified in *A. speltoides DRF1* gene (88% identity between 5’ and 3’ TIRs) through the two sequences alignment (position 710^th^ bp), characterized by 2 mismatches. The presence of the same 2bp TSD and 4bp ITSD was also observed. Furthermore, a 32bp inverted repeat was identified in *A. speltoides DRF1* (90% identity between 5’ and 3’ TIRs) with 3 mismatches and one deletion. Visually, it was also noticed the presence of the same 18bp repeat within the 32bp repeat (Fig. 3), that shared 100% homology with 18bp nested in the 32bp TIR at 5’ end, while at 3’ showed a deletion and a mismatch. A 93bp fragment separated the 18bp TIR and ∼32bp TIR at 5’ end, while an 89bp fragment separated the 18bp TIR and ∼32bp TIR at 3’ end. A 4bp TSD (ACTG) was manually assessed for this 32bpTIR, characterized by 1 mismatch. Presence of ITSD found with 18bp TIR, but no ITSD sequences were found with 32 TIR (Fig. 3). The mismatches due to several mutations are clearly shown in Figure 6. The same 32bp TIR was also manually identified in the *Triticum durum DRF1* gene, but many mutations were present at the 3’ TIR, possibly acquired after domestication of *Triticum durum*.

**Figure 3.**
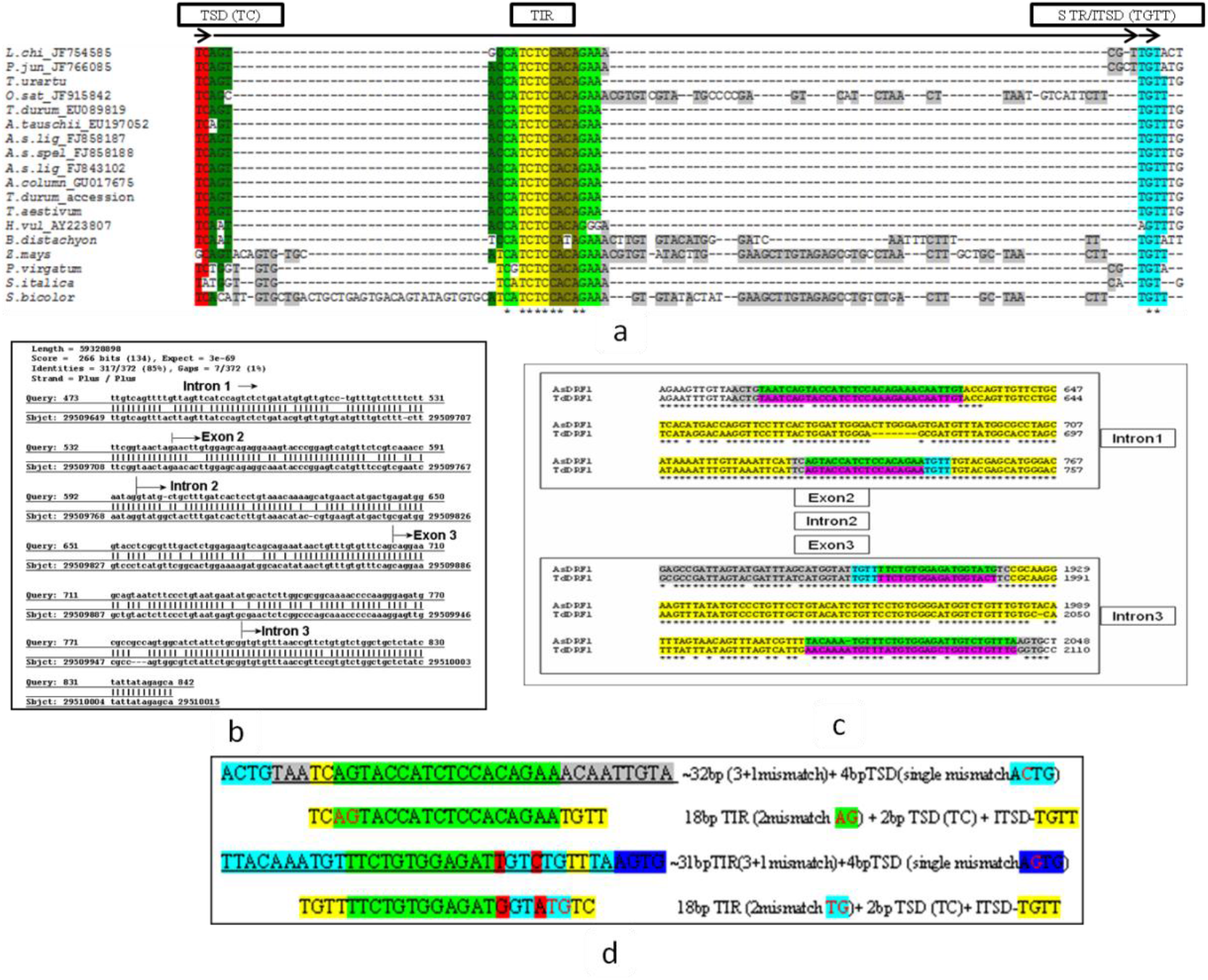
Disruption of 5’ TIR in *Poaceae* members other than *Triticeae* (a), Blast search in the *Brachypodium* genome using the DRF1 transposon sequence as query (b) Alignment showing the variations at 5’ and 3’ on TIR in *AsDRF1* and *TdDRF1* DNA transposons (c) detailed comparison between 18bp and 32bp TIRs in AsDRF1 (d).

All above identified elements, represented the standard transposon signals, thus pointing out the presence of a transposable element in the *DRF1* gene. Apart from Triticeae, the TIR has extremely disrupted mostly through insertion mutations rather than substitutions in other species of Poaceae. Due to insertion the region of TSD, TIR, STR expansion in *S. bicolor* observed with the range of 103 bp, while in *Z. mays* and *O. sativa* the size was 82 bp and 66 bp respectively. The model crop *B. distachyon* found to have the expansion of the TIR with the range of 55 bp. The TIR disrupting insertions were observed before (at 5’) and after (at 3’) the highly conserved region CATCTCCACAGAA in other species apart from *Triticeae*. In all the species within wheat family, the TIR is well conserved. In addition, *P. juncea* was observed with the same, while in *L. chinensis* two substitutions from each region of TIR and STR were observed (Fig. 2). *Z. mays* and *S. bicolor* are closely related species belonging to *Andropogoneae* tribe. The comparative analysis revealed the disruption of (24bp ie TSD; TIR; STR) has revealed well, upon the alignment of this 24bp, used as query. The 24bp sequence has disrupted by insertion and deletions, but not with substitutions due to the observation of significant identify in bases. The alignment resulted with the identity of 95% with *Z. mays* and 91% with *S. bicolor*, at the position of 790 bp distance and 792bp distance relative to ATG respectively in both species. Comparison of this transposable element among different species of Poaceae suggested that during evolution and divergence of species from common ancestor, the recently evolved species in Triticeae tribe including *T. aestivum* was able to retain the footprints of transposon such as TIR, TIR, STR or ITSD etc, while species apart from *Triticeae*, including *B. distacyon* and other species belongs to PACMAD clade were lost their footprints of transposon as indicated in Fig. 2. In contrast, all species were able to retain the core part (Exon 2 and Exon 3) of this transposable element without undergoing any deleterious mutations.

The transposon encompassed two small, transcribed regions (Exon 2 and Exon 3) that were combined by alternative splicing during gene expression. As suggested by CpG plot analysis, the transposon was inserted in a poor CG (about 40%) region (Fig. 4, Panel A). Furthermore, beside all above signals, the core of the transposon (Exon 2 + Intron 2 + Exon 3 and few bps of Intron 1 and Intron 3, at flanking sides) appeared to be very similar to a sequence from *Brachypodium distachyon*, a model grass species. This result appeared very relevant, because it strongly suggested that these non-autonomous DNA transposons may have played a vital role in moving Exon 2 and Exon 3 in grass species during evolution. Probably, it maintained the TIR footprint in most of Triticeae species, but in others, more distant grass species, (as *Brachypodium distachyon*) it was no more present. BLASTn analysis against the whole genome of *B. distachyon* results strongly revealed the novelty, since the presence of the core portion of the transposon without the whole transposon in *B. distachyon*. But there is no evidence for the presence intact transposon in any species prior to the evolution of Triticeae members, thus there is strong evidence of an existence of core element (exon2-intron2-exon3) in species of Panicoideae, Andropogoneae, Brachypoidiae and Oryzeae, however transposon signals and intact transposon carrying the core element is presumably reconstructed during evolution of Triticeae only. Moreover, the undisturbed intact transposon with its core element is present in Chromosome 1A (TGACv1_scaffold_002432 1AL: positions 384010757-384009244, ENSEMBL genomic location: 391258510 bp -391259647 bp), but slightly altered or mutated element is found in Chromosome 1DL (TGACv1_scaffold_062801_1DL) and Chromosome 1BL (TGACv1_scaffold_031713_1BL). However, chromosome 2BL exhibited the presence of partial transposon (IWGSC_V3_chr2BL_scaffold_3259: positions 22051: 22051). Further ENSEMBL wheat BLAST revealed the presence of core element without intact transposon or remnants of transposon in various locations associated with DREB and related genes on Group 1 and Group 2 chromosomes.

**Figure 4.**
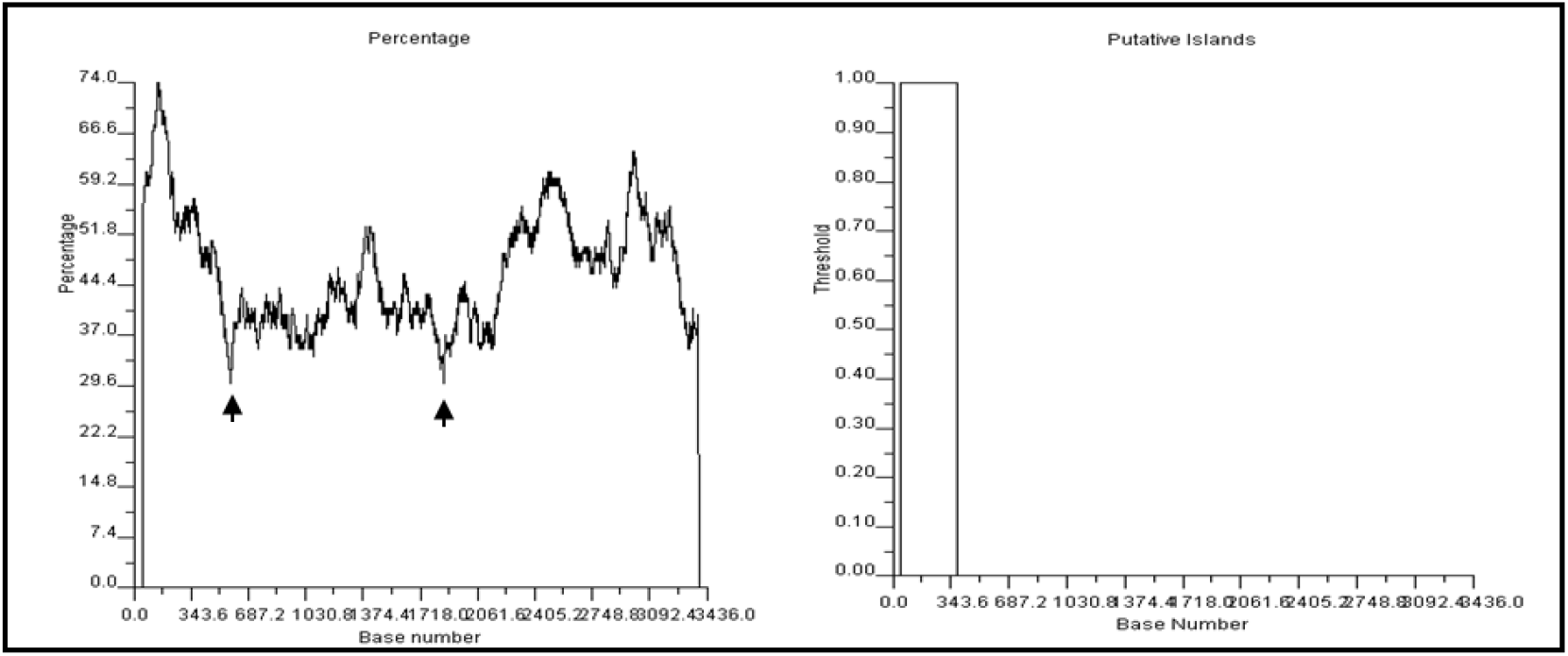
CpGplot results. Panel A: Plot of the percentage of CpG content along the gene sequence. Minimum percentages, located within Intron 1 and Intron 3, possible location of a transposon, are indicated by arrows. Panel B: Putative CpG islands, located within Exon 1.

### Comparison of *Arabidopsis DREB2A* and *DRF1* genes

Comparative genomic study revealed that *DRF1* gene in Poaceae members apart from other monocots had gained additional two exons through the non-autonomous DNA transposon core element inside the gene. While there is no trace or evidence in dicots about the insertion of this transposon mediated core element and the gene is not as large as monocots. BLASTn and ORF analysis revealed that the single exon of dicots corresponds to the last exon (fourth exon) of monocots that codifies for the AP2 domain. Either the whole gene (alternatively spliced) or single exon-based gene is also present in poaceae and can be expressed, which revealed through cDNA search especially in bread wheat. For instance, *DRF1* gene is orthologous/homologous to *DREB2* gene of *Arabidopsis*. During the divergence of mesangiosperm lineage, the gene has been modified drastically. It is known that monocots and dicots diverged from mesangiosperm lineage during early evolution of flowering plants (Zeng et al., 2014), hence the complex form of *DRF1* gene with alternative splicing is observed in poaceae of monocots, especially in triticeae, where the DRF1 gene might have reconstructed with transposon and its core element due to extreme conservation of this transposon and core element particularly in wheat and related species, while there is no any traces of this transposon in dicots and other monocots. Although these orthologus genes sharing similarities and common features in UTR. The alignment revealed that part of the UTR from *DREB2A* in *A*.*thaliana* sharing significant similarity with partial exon1 part of the *DRF1* gene in wheat and its close relatives. Exon1 of *Arabidopsis* is equal to Exon4 of *DRF1* gene, and the only intron of *A. thaliana* has more homology to DRF1 transposon core region encompassing exon2 intron 2 exon3. The protein alignment between *AtDREB2A* and *AsDRF1*.*1* revealed the insertion of small peptide consists of 47 additional amino acids in *AsDRF1*.*1*, while *AtDREB2A* lacks these amino acids due to the absence of transposon mediated coding sequence insertion. Further alignment with *AsDRF1*.*3*, which lacks transposon mediated protein sequence showed better alignment with *AtDREB2A*. In both cases the AP2 domain sequences are highly similar and 84.48 % amino acids are identical among core 58 amino acids of AP2 domain. This indicates apart from transposon region, the key amino acids involving in the interaction with DNA for transcriptional activation, are highly conserved. The AP2 domain is located at exon 4 of *AsDRF1* gene and in the only exon of *AtDREB2A* protein. The calculations of physico-chemical properties of both AtDREB2A and AsDRF1.3 were comparable due to the close vicinity each other. The pI in *AtDREB2A* and *AsDRF1*.*3* (Isoelectric focusing point) was 5.17 and 4.86 respectively and the presence of negatively charged amino acids (Asp+Glu) were 56 and 54 respectively, charged positively amino acids (Arg+Lys) 46 and 35 respectively. Besides, the instability index (II) was computed as 48.59 and 61.91 for *AtDREB2A* and *AsDRF1*.*3* respectively, indicates that these proteins are both unstable.

### Comparison of *TdDRF1* gene sequence with eukaryote repeats with CENSOR server

From 151bp to 230bp, including the junction between Exon 1and Intron 1, a similarity was found with LINE1-42_ZM sequence, a non-LTR Retrotransposon in *Zea mays* whose total length is 6520 bp. Even if they share similarity for just 80bp, perhaps this indicated a fragment from a deleted derivative of the retrotransposon in this part of Exon1 and Intron 1. A second fragment, from 245bp to 294bp, located in Intron 1, showed similarity with hATm-3 AA, from a young family of hATm autonomous DNA transposons, identified in the mosquito (*Aedes aegypty)* genome. A third fragment, from 954bp to 1036bp, also located in Intron 1, showed similarity with DNA-2-16 DR, a non-autonomous DNA transposon from zebrafish. It is worth of noting that this fragment included the tandem repeats of *TdDRF1* gene (Supplementary Fig. 1). The last fragment, from 2671bp to 2718bp, showed similarity with ERV2-1B-LTR_BT, a long terminal repeats from domestic cattle.

### Secondary structures of the *DRF1* transposon

The folding of the *DRF1* transposon was predicted using MFold. The results concerning *Triticum durum DRF1* transposon are shown in Supplementary Fig. 2a. Just two structures were predicted. The same analysis was carried out using *Aegilops speltoides DRF1* transposon and results revealed a more complex picture (Supplementary Fig. 2b). Ten structures were predicted: free energy ranging from -114.31 Kcal/mol to -108.95 Kcal/mol. The first two more stable structures were very similar to each other and comparable to the *Triticum durum* ones, but from the third on, the structures became more complicated, except for the sixth and seventh ones, that were very similar to the first two ones, even if were a bit less stable. The most stable (panel A), an intermediate level (Panel B) and the less stable (Panel C) are shown in Supplementary Fig. 2b. The sequences of the two transposons are quite similar, even if *AsDRF1* transposon contained more repetitions than *TdDRF1* one (e.g. the two copies of 164bp tandem repeat). Possibly, the more complex folds available for *AsDRF1* transposon were related to the presence of more repetitions.

Hypothetical mechanism of TIR, TSD formation and transposon integration is available at Repbase Report server: https://www.girinst.org/server/publ/AsDRF1/ and showed here with sequence (Supplementary Figure 2).

Nonetheless, it is not clear about the hypothesis of the insertion, but it is the most possible mechanism of the intact DRF1 transposon formation due to the presence of strong transposon signals such as Target Site Duplication with Terminal Inverted Repeats. We would also like to mention the noteworthy point that the some species in Poaceae evolved before the evolution of Triticeae members, such as *O. sativa, B. ditachyon, Z. may* didn’t show intact *DRF1* transposon, but showed the intact core element. It indicates that in order to preserve the core element the gene might have modified and reconstructed with non-autonomous DNA during the evolution of wheat and related Triticeae members.

### Divergence time

Lesser disruption of transposon region and TIRs in Triticeae members revealed the recent divergence of this gene in Triticeae members compare to other tribe members in Poaceae. Apart from Triticeae members, species other tribes evolved after triticeae members showed extreme disruptions, especially insertion of additional nucleotides in TIR region and lost the transposon related signatures due to earlier divergence time. The divergence time concerning this transposon sequence from *T. aestivum* to *B. distachyon* and *H. vulgare* was 14 and 4 million years respectively.

The divergence time of *Psathyrostachys juncea* to wheat and its close relatives is about 3 million years. The time of divergence and evolution of bread and durum wheat was very nascent of about 0.02 million years. Comparatively due to recent divergence of wheat and its relatives revealed the lesser disruption or lack of disruption of TIRs/TSDs and intact transposon within Triticeae tribe compared to other Poaceae members in other tribes. Divergence times are consistent with previous studies (to be elaborated in discussion).

## DISCUSSION

This transposon is enriched with tandem repeats, generally, three main types of tandem repeats are reported, namely microsatellites (1-5 bp repetition), minisatellites (6-50bp repetition) and large tandem repeats (greater than 50 bp repetition). Unequal recombination is a major factor for the prevalence of highly repeated sequences in DNA (Smith, 1976). The presence of core element of the transposon or remnants of transposons including tandem repeats are hot spots of recombination (Hellman *et al*., 1988) and cause unusual physical structure and polymerase slippage and result in amplification of many copies core element or remnants of this transposon and its related gene *DRF1* as revealed with our BLASTN analysis from IWGSC, ENSEBL and other servers. It has been shown that the distribution of some transposon families varies with G+C content across the genome (Lander *et al*., 2001). Generally, the transposable element is inserted in a low GC content region or in a region rich A+T, even if many insertions were observed at random site. The most abundant types of MITEs elements, “TA, TAA, and ATA”, *Basho* (T), and *Mariner*-*l*ike *e*lements (TA) are preferably found in A+ T rich regions (Le *et al*., 2000). A putative CpG island was found 50bp to 385 bp within Exon 1 (the observed and expected ratio is greater than 0.60). Other rich regions, with a percentage of CpG content up to 74% were also present. But it is consistent with the insertion of this transposon in target site “TC” with the region of transposon preferred low GC and high AT regions of the gene.

### The transposon and the gene regulation: a possible relationship

We have proposed hypothesis about the insertion of the transposon in *DRF1* gene as available in https://www.girinst.org/server/publ/AsDRF1/. However, this transposon signals were lost and only preserved the core element in tribes evolved earlier than Triticeae, but later evolved wheat and its wild relatives showed an intact transposon with its core element and strong transposon signals. Therefore, it is presumed that the construction or reconstruction of TIR signals around the core element should have presumably happened only in Triticeae members, particularly in wheat and its close relatives. In theory, when a transposon moves, there is the possibility that the included transcribed regions will also move to the target destination in the genome and the continuous transposition could affect the whole gene network system. Thus, since the core of the *DRF1* transposon includes Exon 2 and Exon 3, a possible involvement of this non-autonomous DNA transposon in the alternative splicing regulation of the *DRF1* gene was investigated through genomic approach. In fact, the transposon could move, excising the two exons from the gene. But the hypothetic resulting 255bp sequence was not present anymore, as assessed by BLAST analysis, suggesting that an unknown mechanism prevents further movements of the *DRF1* transposon. Methylation of transposons has been reported as a natural epigenetic phenomenon resembling transgene silencing for the defence of genome (Yoder *et al*., 1997; Matzke *et al*., 1999; Selker *et al*., 2003).

Transposon regulation is tuned along the sequences of heterochromatin regions in plant and animals and is also controlled by RNAi (Lippman *et al*., 2004). Besides the methylation protection mechanism, the activity of *DRF1* transposon could be suppressed by RNAi. Leeds *et al*., (1991) identified the *UPF* gene as the responsible for the degradation of mRNAs containing premature stop codons (PTC). They demonstrated that UPF gene does not affect the normal mRNAs of *HIS4*/*LEU2* genes but affects *HIS4*/*LEU2* mRNAs with PTC. The *UPF* gene is widely present in most of the eukaryote species, but not in prokaryotes and Archeaebacteria (Culbertson *et al*, 2003). Another factor, namely an unknown endonuclease, involved in the degradation of mRNAs with premature stop codons in *Drosophila* was described by Valencia-Sanchez and Maquat (2004). Baulcombe (2005) suggested that normally mRNAs are targeted by exonucleases and subsequently the aberrant mRNAs are targeted by RNAi, due to lack 5’of cap and/or PolyA tail. Really mRNAs containing PTC are present among the transcripts of *DRF1* gene. The *DRF1* gene, due to alternative splicing, produces three mRNAs, one of them, *DRF1*.2, containing a PTC. It was demonstrated that *DRF1*.2 is constitutively produced at large quantity (Latini *et al*., 2007). Thus, it should be degraded due to the presence of PTC and the aberrant RNA can be targeted by RNAi, triggering the nonsense mediated decay (NMD) regulation pathway, or can trigger the silencing of the transposon (Zamore, 2002). *DRF1* organization consists of four exons that combine to obtain the three transcripts. *TdDRF1*.*1* consists of all exons producing the largest transcript, about 393 amino acids long. *TdDRF1*.*2* consists of Exon1, 2 and 4, the junction of Exon 2 and 4 resulting in the formation of a PCT. *TdDRF1*.*3* consists of Exon 1 and 4 and produces a transcript about 346 long. Thus, when both the exons inside the transposon (Exon 2 and 3) are present (*DRF1*.*1*) or both are absent (*DRF1*.3), the resulting transcripts do not undergo non-sense mediated decay because PTC is not present. On the other side, the presence of only one of them, namely Exon 2, directly combined with Exon 4 (*DRF1*.2), promotes the degradation of this transcript, resulting in the formation of PTC and finally NMD (Fig.6). The complexity of the *DRF1* gene structure suggested further investigations. The NetGene2 server was used to assess the exon intron splice sites and branch points in the *DRF1* gene. Besides the expected ones, in Intron 3 two branch points were found, an unexpected donor splice site and one unexpected acceptor splice site. Branch point sequences in introns can become core signals for splicing. Thus, Intron 3 can be considered as a pseudo exon and further investigations were carried out. Initially, the transposon sequence was manually split in its components and the fragments used as templates in BLAST analysis. Because results concerning Intron 3 fragment appeared interesting, its whole sequence was also analysed. Hence a BLAST search at TIGR Wheat Genome Annotation database was carried out using Intron 3 sequence as query. The best result using *Triticum aestivum* release 2 database concerned the expressed sequence BQ240043, mRNA obtained from wheat seeds of cultivar Glenlea 5 days post-anthesis, two long fragments aligned showing 74% and 83% identity, E-value= 1.8 e-72. Other relevant results concerned three sequences, namely CJ729917, BE637924 and CJ630958, all related to DREB transcription factor 3A. When the search was carried out against DFCI wheat gene index, a large contig TC263997 was retrieved, that included a part of BQ240043. A BLASTn analysis carried out in GrainGenes retrieved more or less the same sequences arranged in different contig. In the case of *TdDRF1*, the sequence of Intron 3 is not expressed, but the above results demonstrated that highly similar sequences were found expressed. It is known that intron regulatory elements, ISE (intronic splicing enhancer), or ISS (intron splicing silencer) enhance or inhibit the use of adjacent splice sites within intron. Transacting splicing factors (SRE-Splicing Regulatory Elements) can activate or suppress the splice site recognition or promoting various mechanisms for spliceosome assembly (Matlin *et al*., 2005; Chasin *et al*., 2007). Thus, it is possible that the sequence, which represents an intron in *TdDRF1* gene, is indeed expressed, in other circumstances. Otherwise, because it is part of a transposon, it is possible that Intron 3 sequence moved towards other target regions and became part of expressed sequences. It is known that intronic or exonic splicing enhancer that activated by splicing machinery with transacting splicing factors (Venables et al., 2011). This theory is further supported by the dentification and analysis of cis element that present in exon 3 called as exonic splicing enhancer (ESE1) in *LcDREB2* (Liu et al., 2017).

**Figure 5.**
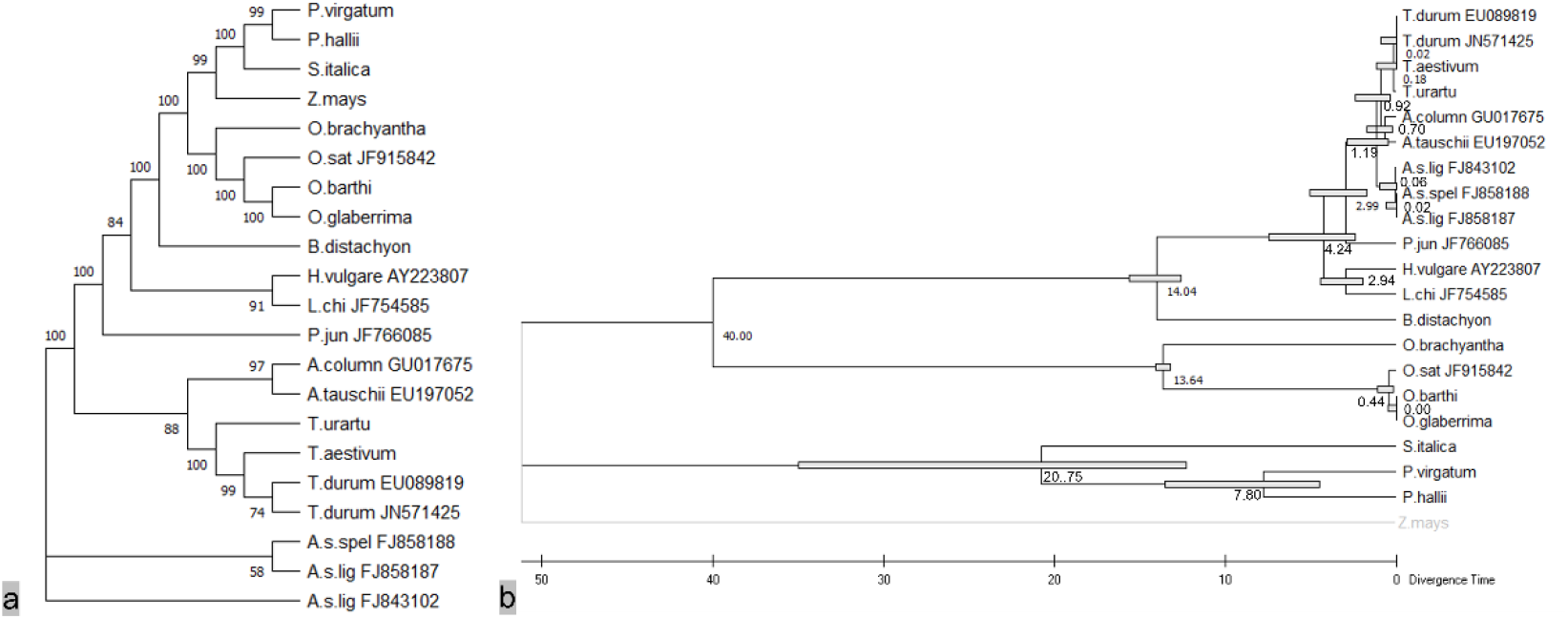
a) Maximum likelihood tree generated with MEGA11.0.8 using diverse Poaceae species with 1000 bootstraps through Tamura and Nei model. b) Divergence times for all branching points was computed using Maximum Likelihood method and Tamura-Nei model. Anchor point was set at the node between *O. sativa* and *T. aestivum* (Divergence time 40 million years) and divergence time estimated using RelTime-ML.

**Figure 6.**
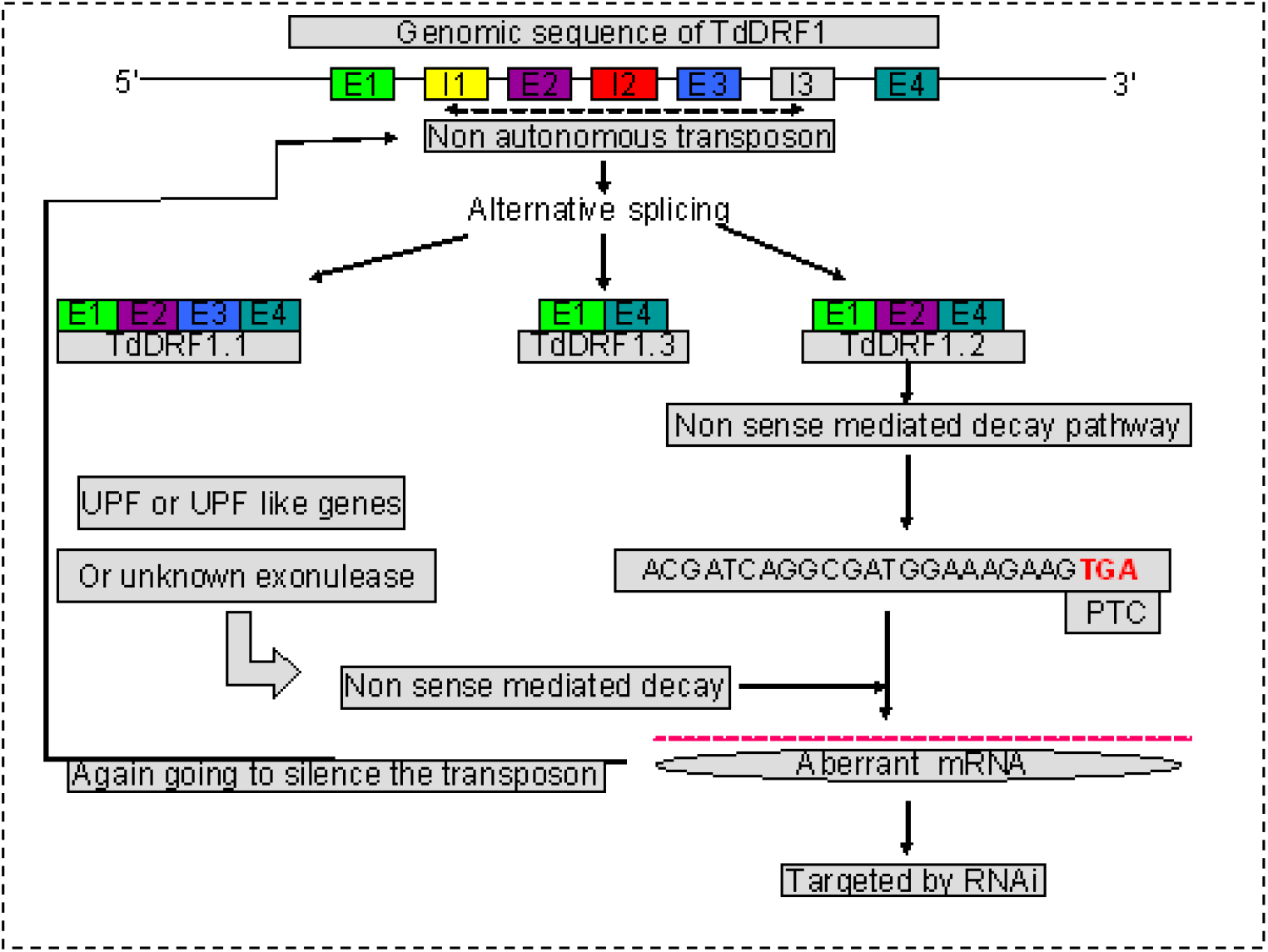
Expression of alternatively spliced transcripts in *DRF1* gene and hypothetical mechanism behind in the suppression of a *DRF1* transposon.

### A comparison of cis acting core elements in DRF1 transposon and alternative splicing

The analysis of *Arabidopsis DREB2* gene (including intron and UTR) with *DRF1* gene and its transposon revealed the sharable features of the gene except the transposon region. *DRF1* is dicot ortholog of *DREB* gene of Monocots, however none of the *DREB* dicot orthologs exhibited the presence of this transposon insertion as revealed by this study and none of the *DREB* genes in *A. thaliana* exhibited alternative splicing. The alternative splicing pattern of DREB gene subgroup 2 is confined to Poaceae within A-2 group (Liu et al., 2017). It is not known why *Z. mays, B. distayon, O*.*sativa* kind of Poaceae members didn’t show the DRF1 transposon signals or showed highly disturbed transposon signals that not expected to be a transposon. However, these species contain absolute core element (exon2-intron2 and exon3). Alternative splicing is firmly regulated by the interaction of trans acting factors on cis elements in exons and surrounding introns (Lee et al., 2015). In *DRF1* gene, it is possible that alternative mode of splicing was even present in the gene even before the evolution of actual intact transposon in *Z. mays, B. distayon, O*.*sativa* kind of organisms. Also, members apart from wheat and its progenitors such as *H. vulgare* and *L. chinensis* were also showed somewhat disturbed transposon or transposon signals. Thus, apart from Poaceae, other monocot members lacking any transposon and its signal in DRF1 or any DREB related gene such as *Asparagus officinalis, Musa acuminata* and *Dioscorea rotundata*. However, Poaceae members especially in Triticeae tribe, wheat and its progenitors only showed an intact transposon and transposon signals. Some questions yet to be answered.

1. Whether complex form of this *DRF1* transposon and its signal lost in DRF1 gene of Poaceae members other than Triticeae tribe during evolution as they lost their transposon signals and only preserved the core element?
2. Why are this transposon and its signals well preserved in wheat and its progenitor alone even not in other Triticeae members such as *H. vulgare* and *L. chinensis*?
3. Whether the nascent evolution of wheat and its progenitors expected to have lesser spontaneous mutations and showed without disturbance of intact transposon and transposon signals?

This transposon harbouring ortholog of monocot sharing some common features with dicot *A. thaliana* such as Exon1 of *DRF1* showed partial identity with 5’ UTR in *Arabidopsis*, Exon1 of *Arabidopsis* is equal to Exon4 of *DRF1*.Only one Intron in *A. thaliana* didn’t show any homology to core transposon region i.e., exon2 intron 2 exon3 part of DRF1 gene. This study further supporting the hypothesis that monocots diverged from dicot relatives during the very earlier course of evolution of flowering plants (Dahlgren and Clifford, 1982, Duvall 1993). Therefore, the *DREB* gene related to *DRF1* gene had simplified form of genetic organization with single exon encoding feature, while the divergence of monocots resulted with reshaping of this gene and in Triticeae particularly in wheat and its progenitors, the DRF1 transposon is well preserved or reconstructed during evolution. Spontaneous changes in DNA such as mutations or other genetic modifications resulted the aberration in transposon signatures in Poaceae members apart from Triticeae. Nevertheless, the core element carried by the transposon is well preserved in almost all Poaceae members, while dicots and other monocots neither exhibited the presence of transposon with alternative mode of splicing nor the core element.

## Conclusion

The analysis of this transposon revealed that the core element and mode of alternative splicing was present even before the evolution of this transposon but only in Poaceae members rather than other monocot and dicot species. However, it is yet not known how this transposon is intact with strong transposon signals in later evolved wheat and close relatives in Triticeae rather than formerly evolved species of other tribes. The presence of core element that contains a cis element according to existing evidence reveals the regulation of alternative splicing is an important evolutionary driving force within Poaceae to increase the gene expression with alternative splice transcripts in *DRF1*, a *DREB* related gene. Nonetheless, the preservative nature of *DRF1* transposon signals and the intact transposon might have constructed or reconstructed during the evolution of Triticeae members particularly in wheat and its close relatives and it is expected to play a major role in evolutionary driving force associated movements of the core element in *DRF1* and *DREB* related genes.

## Declaration

The authors declare no competing interests.

---

**Supplementary Figure 1.**
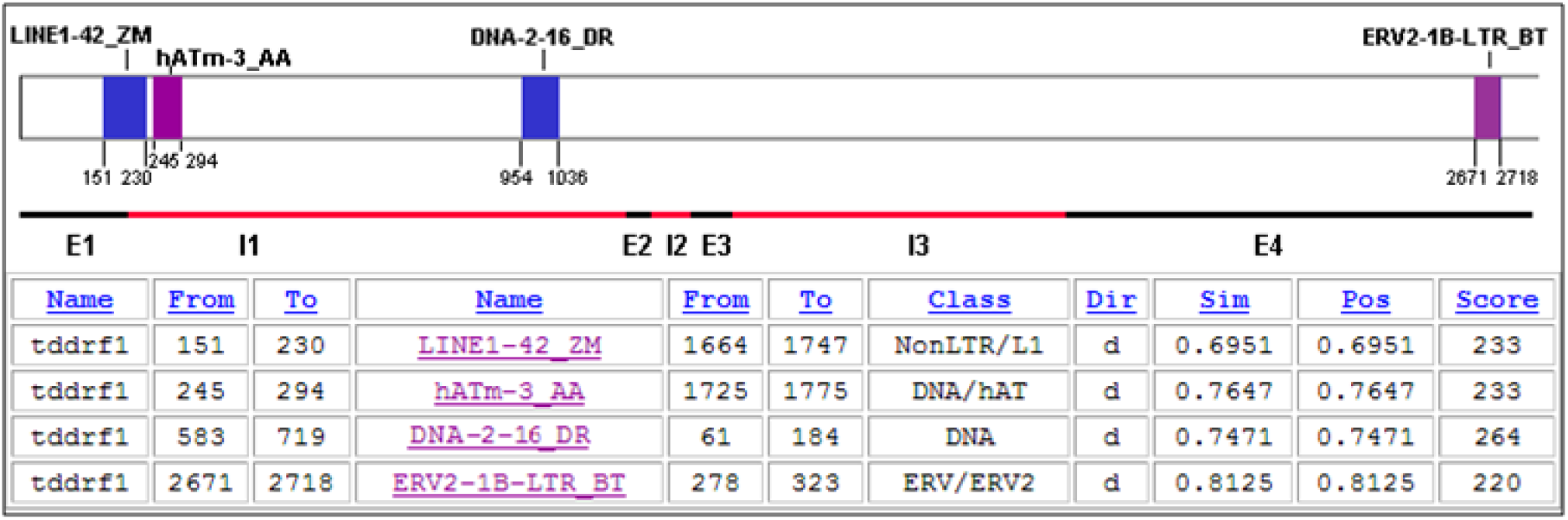
Repbase-CENSOR analysis of the repeats in *TdDRF1* gene.

**Supplementary Figure 2.**
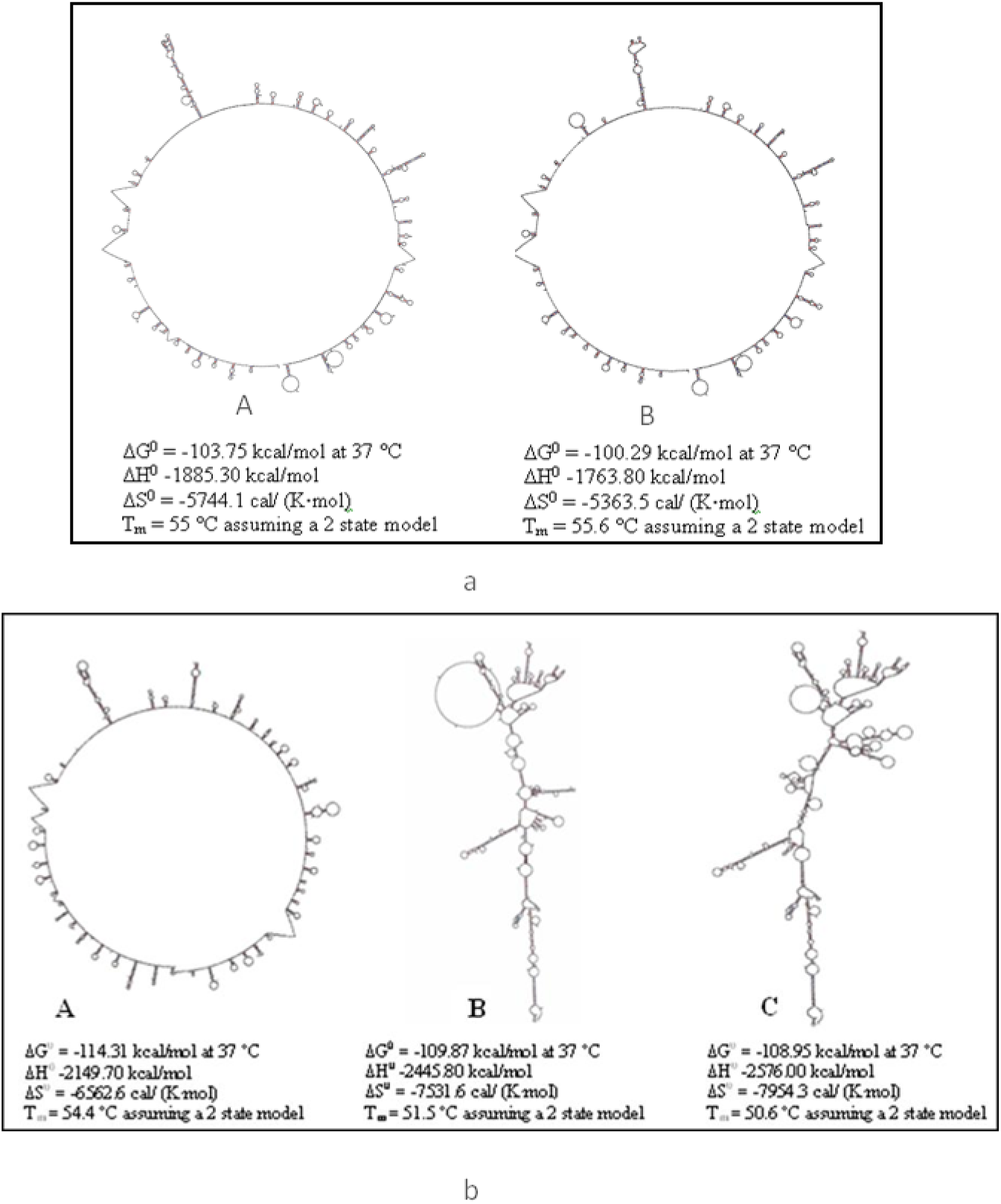
a) Two structures calculated for *TdDRF1* transposon, b) Three of the ten structures calculated for *AsDRF1* transposon

**Supplementary Figure 3.**
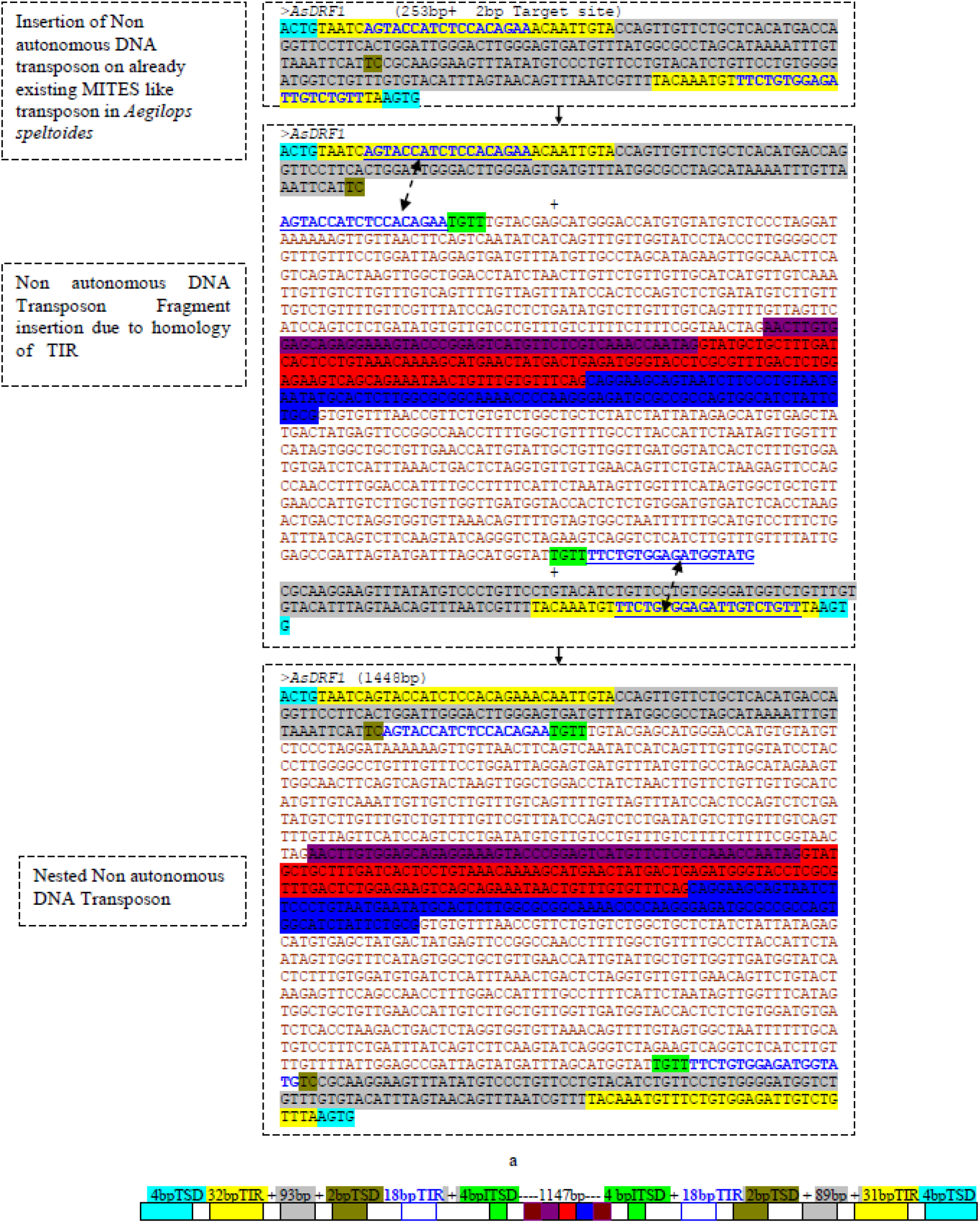
a) A possible mechanism of *DRF1* transposon insertion carrying two exons and an intron, b) Final length of the nested non autonomous *AsDRF1* Transposon 1448bp.

## Notes

https://www.girinst.org/2009/vol9/issue2/TdDRF1.html

https://www.girinst.org/2009/vol9/issue3/AsDRF1.html

http://www.girinst.org/server/publ/AsDRF1/

https://om.ciheam.org/om/pdf/a110/00007124.pdf

### Competing Interest Statement

The authors have declared no competing interest.

